# Evolutionarily diverse origins of honey bee deformed wing viruses

**DOI:** 10.1101/2023.01.21.525007

**Authors:** Nonno Hasegawa, Maeva A. Techer, Noureddine Adjlane, Muntasser Sabah al-Hissnawi, Karina Antúnez, Alexis Beaurepaire, Krisztina Christmon, Helene Delatte, Usman H. Dukku, Nurit Eliash, Mogbel A. A. El-Niweiri, Olivier Esnault, Jay D. Evans, Nizar J. Haddad, Barbara Locke, Irene Muñoz, Grégoire Noël, Delphine Panziera, John M. K. Roberts, Pilar De la Rúa, Mohamed A. Shebl, Zoran Stanimirovic, David A. Rasmussen, Alexander S. Mikheyev

## Abstract

Novel transmission routes can allow infectious diseases to spread, often with devastating consequences. Ectoparasitic varroa mites vector a diversity of RNA viruses and, having switched hosts from the eastern to western honey bees (*Apis cerana* to *Apis mellifera*). They provide an opportunity to explore how novel transmission routes shape disease epidemiology. As the principal driver of the spread of Deformed Wing Viruses (mainly DWV-A and DWV-B), varroa infestation has also driven global honey bee health declines. The more virulent DWV-B strain has been replacing the original DWV-A strain in many regions over the past two decades. Yet, how these viruses originated and spread remains poorly understood. Here we use a phylogeographic analysis based on whole genome data to reconstruct the origins and demography of DWV spread. We found that, rather than reemerging in western honey bees after varroa switched hosts, as suggested by previous work, DWV-A most likely originated in Asia and spread in the mid-20^th^ century. It also showed a massive population size expansion following the varroa host switch. By contrast, DWV-B was most likely acquired more recently from a source outside Asia, and appears absent from eastern honey bees, the original varroa host. These results highlight the dynamic nature of viral adaptation, whereby a vector’s host switch can give rise to competing and increasingly virulent disease pandemics. The evolutionary novelty and rapid global spread of these host-virus interactions, together with observed spillover into other species, illustrate how increasing globalisation poses urgent threats to biodiversity and food security.

## Introduction

Increased globalisation, trade, and habitat fragmentation bring previously isolated species into contact, which risks increased transmission of diseases and parasites. These new connections caused global pandemics affecting populations of humans, animals and plants. Many of these outbreaks have been caused by RNA viruses, and studies have highlighted them as primary etiological agents of human disease (1–4). The high adaptive potential of RNA viruses is linked to their high mutation rates, which are often an order of magnitude higher than those of DNA viruses (5) and their host range is frequently limited by their capacity to reach novel host populations (5, 6). Yet, our ability to predict specific viral host shifts and their consequences remains poor.

Many viruses solve the challenges of transmissibility by using vectors to connect potential hosts. As a result, changes in vector ecology, such as range expansions or host shifts can drive viral pandemics. One model of viral spread involves anthropogenic changes in vector range, allowing viruses to spread to new geographical areas. For example, climate change has led to the poleward expansion of mosquito ranges bringing dengue to new human populations (7). Alternatively, viruses can acquire a novel host, driving their spread. This has happened in the case of the Chikungunya virus, originally found in arboreal mosquitoes in primates in sub-Saharan Africa, but has switched to urban-dwelling mosquitoes and spread through the human population to other continents (8). While human viruses have been studied extensively, these ecological and evolutionary dynamics are not limited to our species. Globalization has also caused pandemics among non-human animals and plants (9, 10). Yet, our ability to follow the dynamics of these pandemics is typically limited by a poor understanding of viral ecology in the original host, if the host is known at all.

Exceptionally, given their importance in world agriculture, there are excellent historical records on the spread of honey bee (genus *Apis*) parasites and viruses (11, 12). With their rapid generation times and large population sizes, arthropods provide an excellent opportunity to examine how vector host switches affect viral evolution. The eastern honey bee (*Apis cerana*) and the western honey bee (*Apis mellifera*) are sister species that diverged ~7 million years ago (13). Eastern honey bees are found throughout Asia, while western honey bees naturally range from Europe to Africa but are nowadays distributed all over the world except Antarctica. Western honey bees have larger colony sizes and produce more honey, making them the mainstay of the beekeeping industry worldwide. Western honey bees existed in allopatry from other congenerics, isolating them from many honey bee diseases and parasites. However, that changed in the 19th century when beekeepers took western honey bees to Asia where they came in contact with eastern honey bees and *Varroa* mites, which are obligate parasites of honey bee brood. By the mid-20th century, *Varroa destructor* switched to its novel honey bee host (*A. mellifera*), followed by *Varroa jacobsoni* in the early 21st century (11, 14, 15). *V. destructor* then spread worldwide, transmitting a wide variety of bee viruses and causing global population collapses of western honey bees (12).

The well-documented history of the varroa host switches makes this an excellent system to examine how novel transmission routes affect viral disease emergence, as well as providing insights into the evolution of viral diseases (16). Here we focused on understanding (1) how host bee species affect the virome of varroa and (2) reconstructing the geographic spread of the two most common and important viruses vectored by varroa – deformed wing virus (DWV) strains A and B that diverged about 200 years ago (17). A previous phylogeographic investigation of DWV-A found that the virus most likely originated in Europe and has increased in frequency after the introduction of varroa mites (18), though the inference was based on less than 10% of the viral genome. The origins of DWV-B remain poorly understood to date, though its spread in honey bees appears to have happened in recent decades (19). While these two strains are similar in sequence and biology, they differ significantly in how they interact with both bees and mites (20, 21). This study aimed at using full-genome sequences to examine whether these viruses had similar or different phylogeographic histories, evolutionary origins and population dynamics.

## Methods

Data analysis pipelines and R scripts are available at: https://github.com/nonnohasegawa/varroa-transcriptomics.

### Mite collection, sequencing and bioinformatic analysis

Varroa mites were collected from 1989 to 2019 from 43 countries and islands spanning six continents (Table S1 and Figure 1A). Mite RNA was extracted from single adult female mites using TRIzol and prepared for sequencing using the previously described methodology (22). Sequencing was performed on an Illumina Novaseq S2 at 150 cycles with paired-end reads. Raw sequence data was first mapped to *Varroa destructor* reference genome Vdes_3.0 [GCF_002443255.1] (23) using Bowtie2 (24). The unmapped reads from the previous step were mapped to a selection of viruses that are known to be associated with varroa mites (21). Using VariantBam (25), duplicates were removed and downsampled to a maximum coverage to 200. A pileup format was created using SAMtools (26), then variants were called using VarScan (27). Indels, which are phylogenetically uninformative, were removed using vcftools (28). Then viral sequences of each sample were extracted using BCFtools (26). In addition, we quantified viral abundances using Kallisto (29) on raw reads mapped to reference viral genomes.

**Figure 1.**
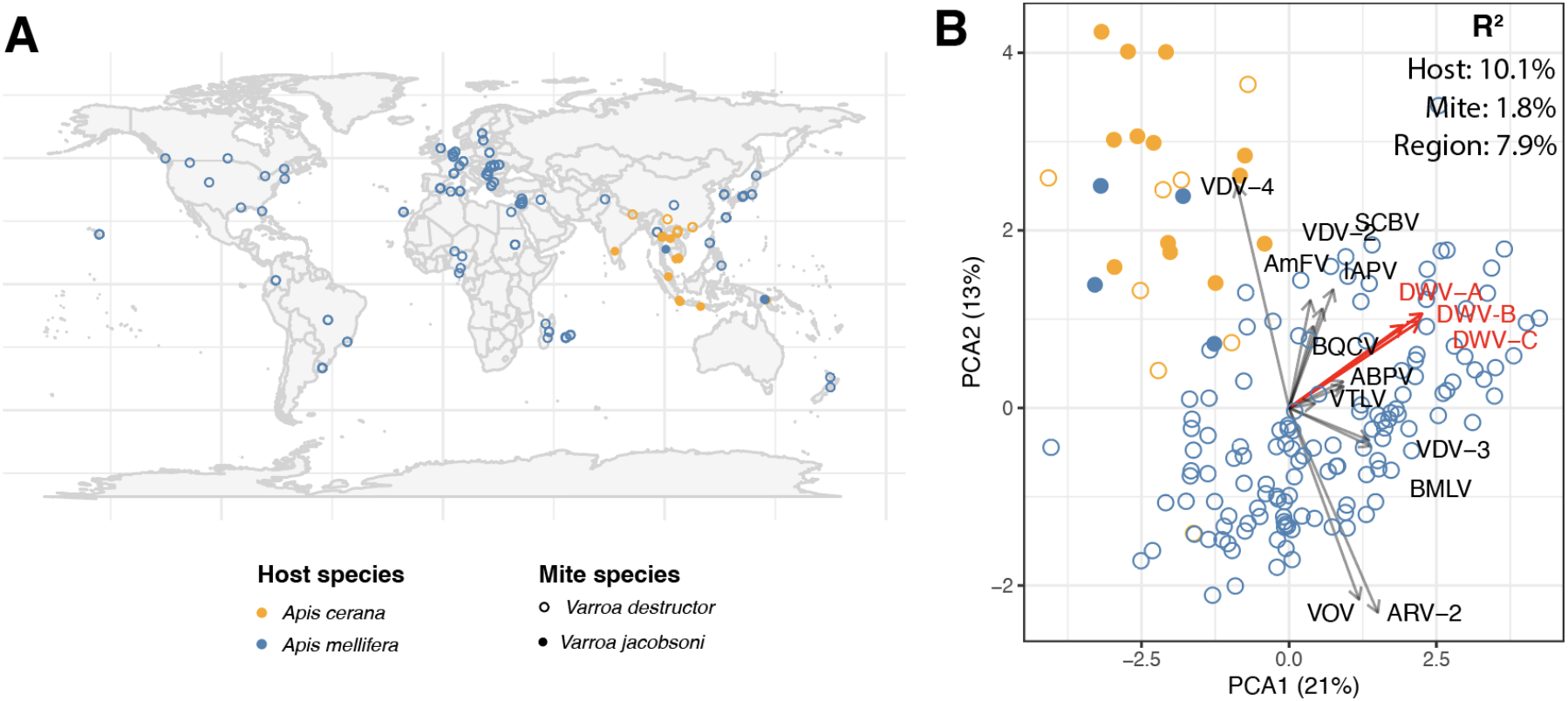
Varroa viromes are most strongly affected by the host bee species. (A) Map of 166 samples used in the virome analysis. (B) The PCA plot of varror virome composition. Arrows represent loadings of each virus onto the principal component axis (*i.e*., how correlated its titre is with each of the two PCA axes). Loadings show that mites from eastern honeybee have generally lower titres of deformed wing viruses (DWV-A, DWV-B and DWV-c), as well as most other viruses, except VDV-4, which was more abundant. These community-level data shows that the viral landscape inside the mite is most likely shaped by the host bee, either by direct uptake of viruses by mites from bees, or via tripartite coevolutionary relationships. This relationship is quantified by R^2^ values from an analysis of variance of viral abundance distance matrices, suggesting that much of the mite virome was acquired from the host, relative to the effects of the mite itself. However, all effects were significant (p < 0.001), indicating that the virome is shaped by all three factors.

### Host, mite species and geographic effects on the virome

For the analysis of virome composition, we first filtered out 8 libraries which had unusually low read counts. Read counts are compositional data, which means that an increase in one virus abundance in a sequenced sample must lead to a proportional decrease in the abundances of other viruses. To address this issue we performed an additive log-ratio transform on the count data (30), using the number of reads mapped to the varroa genome as a common denominator. The resulting ratios should be independent of each other. We then used the ratios to perform a principal component analysis and to quantitatively assess the effects of honey bee host species, mite species and geographic region using analysis of matrix distances analysis using Euclidean distances (31). Statistical analyses were conducted on a total of 166 libraries using the R programming environment using the `vegan` package (32).

### Phylogeographic analysis

Consensus sequences extracted from varroa mite transcriptomes were aligned. We excluded genomes with insufficient sequence coverage, less than 90% for DWV-A and 80% for DWV-B. To this dataset we added complete viral genomes of DWV-A and DWV-B from NCBI and consensus sequences from a published dataset (33), excluding those identified as recombinants by other studies (Table S2). In order to improve alignment quality, all genomes were trimmed to include just the polyprotein and codon aligned. All viral genomes were screened for possible recombination events using RDP4 (34). Sequences were considered likely recombinants if most detection methods implemented in RDP4 showed significant evidence for recombination. Overall we identified one DWV-A sequence (out of 56) and two DWV-B sequences (out of 32) that were likely to be recombinants and excluded them from further analysis that relied upon a tree-like sequence topology.

The phylogenetic history of DWV isolates was reconstructed jointly with their geographic movement using a discrete-trait CTMC phylogeographic model in BEAST v. 2.6 (35, 36). Viral sequences were assumed to evolve under a General Time Reversible substitution model with Gamma-distributed rate heterogeneity across sites. To obtain a time-calibrated tree, we assumed a strict molecular clock with a LogNormal (mean=0.001;sd=2.5) prior to the clock rate. For the phylogeographic reconstruction, sampling locations were binned into seven discrete locations (Africa, Asia, Europe, the Mediterranean, North America, Oceania, and South America). Bayesian Stochastic Search Variable Selection was used to identify movement rates between locations significantly higher than zero (36). The geographic origin of each DWV strain was then reconstructed based on the (posterior) ancestral state/location probabilities computed at the node representing the MRCA of each strain. Inferred origins, therefore, take into account uncertainty both in the phylogenetic (tree) and reconstructed ancestral states. We provide the input BEAST XML files for further details on the phylogenetic analysis (Supplementary Dataset 1).

We also reconstructed the demographic history of each DWV strain using a Bayesian Skyline analysis in BEAST 2 (37). To account for phylogenetic uncertainty, phylogenetic trees were jointly reconstructed along with demographic parameters using the same substitution and clock models as above. Historical changes in effective population sizes (*N_e_*) were reconstructed under a Skyline coalescent tree prior 10 coalescent time intervals, each with a Jeffrey’s prior on *N_e_*.

## Results

Each library generated an average of 9,216,119 reads (s.d. 3,057,629), of which an average of 57.7 % (s.d. 12.8%) mapped to viruses (ranging between 21.7 – 93.3%). We extracted 83 viral genome sequences, including 54 from our study adding a further 29 reference sequences for phylogenetic analysis (Table S1 and S2).

### Host effects on the virome

Our data set consisted of RNA seq data that captured viruses from two species of mites collected from either original or novel bee hosts (*A. cerana* and *A. mellifera*). We used these data to determine the relative importance of host and vector species for the composition of the varroa virome, which showed that the host bee species played a greater role than the species of the mite (Figure 1). Viromes of *V. destructor* mites which were collected from *A. mellifera* host had a higher number of viruses present, notably higher numbers of deformed wing viruses (Figure 1B). On the contrary, viromes of varroa mites collected from *A. cerana* hosts exhibited higher association with VDV-4 and not to deformed wing viruses. We had far fewer *V. jacobsoni* samples, but they appeared to show a qualitatively similar trend (Figure 1B).

### Origins of deformed wing viruses

We found that DWV-A and DWV-B diverged from each other 308 years ago (95% HPD 266 - 353 years), predating the varroa host switches. There are other DWV strains, the most common being DWV-C. While we did not encounter enough of this strain to conduct phylogeographic analysis, we could compute its divergence from the other two strains, which was even more ancient, 979 years ago (HPD 836 - 1128). We found strong evidence that DWV-A originated in Asia (Figure 2A), and that its population expanded rapidly after the varroa host switch to western honey bees, consistent with spillover onto a novel host (posterior probability = 0.97). By contrast, while the most ancestral DWV-B samples originate from the Middle East, this origin was less supported (posterior probability = 0.48), and no evidence of an increase in post-varroa host switch population size could be detected (Figure 2B). However, an Asian origin for DWV-B is not likely (posterior probability = 0.12).

**Figure 2.**
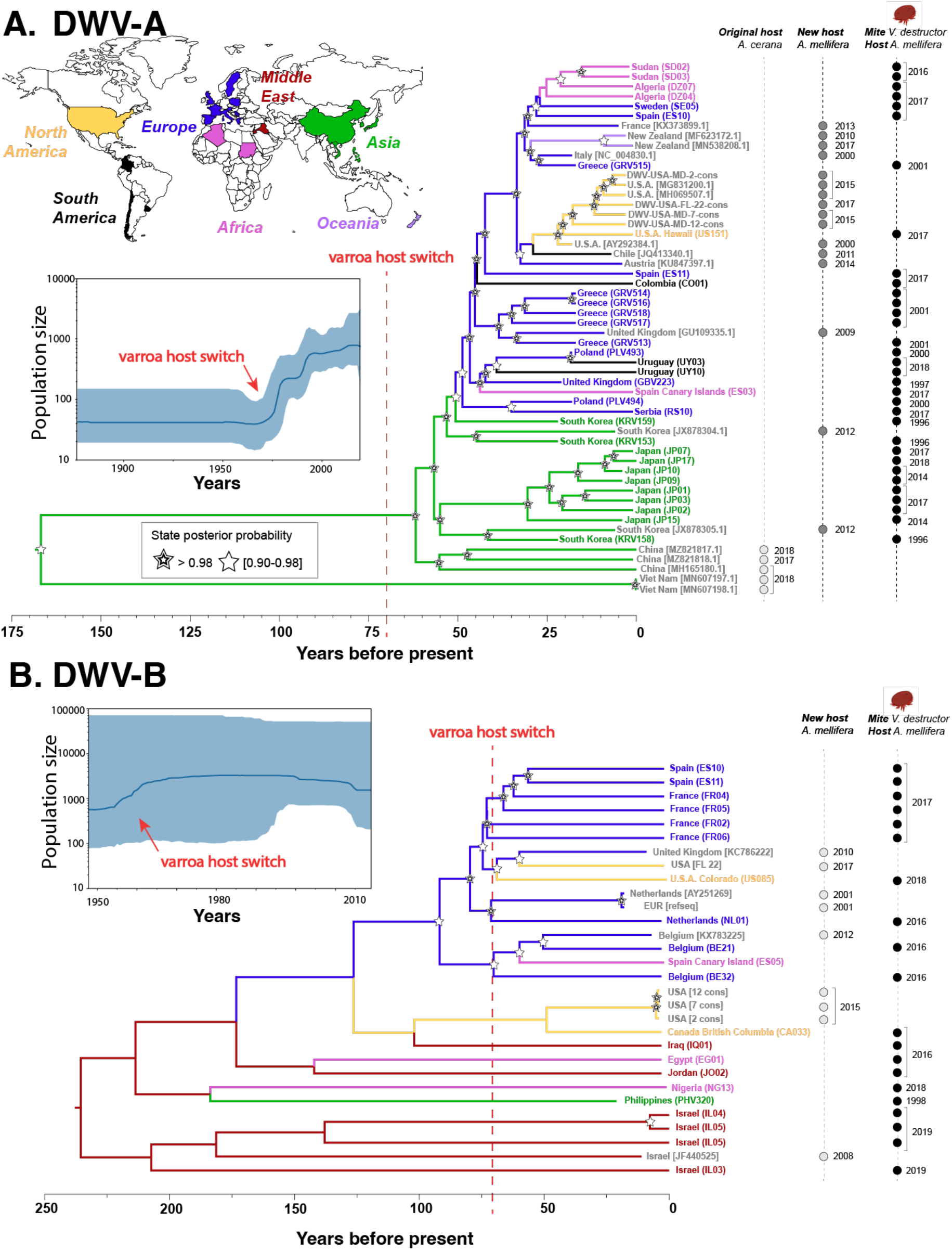
Divergence and phylogeography of deformed wing viruses based on sequences from mites and bees. Locations, where samples were collected are colour-coded. Sample identifiers refer to samples sequenced in our study (Table S1) or via their NCBI identifiers (Table S2). (A) Divergences between the three main DWV strains predate the host switch by *V. destructor* by hundreds of years. Phylogeographic reconstruction indicates that DWV-A originated in Asia (posterior probability 0.97). (B) By contrast, the provenance of DWV-B remains unclear with the strongest evidence for a Middle Eastern origin (posterior probability 0.48). Bayesian skyline reconstruction shows a sharp increase in DWV-A populations in the mid-20th century concurrent with varroa’s host switch and subsequent global spread. This suggests that the host switch was mechanistically linked to the demographic expansion of the virus. By contrast, no such pattern could be seen in DWV-B, which appears to have a much more stable population size, although there is strong evidence that it out-competes DWV-A in honey bee colonies (19, 33, 41). Also, note the different time scales in the two panels, with most divergences in the DWV-B tree being much deeper than those in DWV-A. These data show that the two closely related viruses most likely have distinctly different geographic and evolutionary origins as well as dynamics of spread, while predominantly relying on the same host and vector.

## Discussion

Host availability limits the spread of many RNA viruses, and changes in vector ecology can lead to novel transmission pathways. For example, climate-driven expansion of mosquito ranges leads to larger endemic ranges of many mosquito-borne diseases (7). Alternatively, acquiring a new host and/or vectors can allow a virus to spread globally, as in the case of the Chikungunya virus (8). Mirroring these human vector-borne viruses, we found evidence for both mechanisms in DWV. DWV-A spilled over into western honey bees in association with the varroa host switch. While the exact geographic origin and host of DWV-B remain unclear, several lines of evidence suggest it was acquired by varroa after its host switch. First, it appears to be absent in eastern honey bees, whereas DWV-A is detected consistently (38–40). Second, its split from DWV-A predates the varroa pandemic by hundreds of years (Figure 2). Third, epidemiological data suggest a spread that started over the past two decades, long after the varroa host switch that started in the mid-20th century (19, 33, 41). These lines of evidence point to the dynamic interplay between vector and disease ecology, and the potential for continuing viral emergence after a vector host switch.

Taking advantage of viromes collected from different species of mites across different bee hosts, we could ask which factors play a greater role in shaping viral community composition. Most of the differentiation was associated with the bee host rather than the mite species (Figure 1). These findings are interesting in light of recent work suggesting that viruses interact with varroa gene expression in species-specific ways that affect viral titres (21). However, relatively few viruses replicate in varroa, and therefore must be taken up from the bee hosts. Thus, the biology of the host bee likely remains the primary driver of viral community composition.

It appears that DWV-A was already present in eastern honey bees and spilled over to western honey bees at the same time as the varroa host switch (Figure 2). These findings differ significantly from earlier work that excluded an Asian origin and suggested that the virus was endemic to western honey bee populations before the arrival of varroa (18). The reason for these discrepancies is not clear, though our study used an order of magnitude more sequence data per viral genome, and could pick up stronger signals, such as the explosive growth of DWV-A associated with varroosis that could not be detected by the previous study (Figure 2A).

Our findings that varroa drove the spread of DWV-A need to be reconciled with other work that found DWV-A in regions where varroa has not yet been detected, for instance as in Australia (18) or in advance of the invasion front in Hawaii (42). There are several possibilities. It is possible that once DWV-A became established in western honey bees post-varroa host switch, it gained access to additional transmission pathways that allowed spread without varroa. It is also virtually certain that some of these results were caused by contamination. *E.g*., a subsequent in-depth study of Australian viruses, where libraries were prepared and sequenced Australia, without the possibility of cross-contamination, failed to find DWV (43). To further illustrate this point, a cursory examination of sequenced negative controls in the NCBI SRA database (search terms `negative control AND “Apis mellifera”[orgn]` in RNA data, 19 January 2023), yielded just two sequencing runs of library blanks (44), though both have DWV contamination present (accession numbers DRR333172 and DRR333191). We recommend that future studies aiming to show the presence of DWV, sequence negative controls as standard practice to rule out contamination. This is particularly important for studies aiming to detect viral spillover into non-*Apis* taxa, which are usually prepared and sequenced alongside honey bee samples.

In contrast to DWV-A, it appears that DWV-B was either present in western honey bees or was acquired from another host after the varroa host switch. It was first detected in 2004 (45), suggesting that, barring extensive underreporting, it was either a novel or geographically restricted virus. While bearing in mind the caveat above regarding contamination, DWV-B is found in other taxa (46, 47), and some of them could be potential sources of infection. Interestingly, despite recorded increases in DWV-B levels in honey bees, we failed to detect an increase in population size from molecular data, in contrast to DWV-A (Figure 2). This could suggest that there is a large alternative host population other than honey bees, or simply that the population size increase has not reached the same levels as observed with DWV-A. Future work should investigate the origins of DWV-B in order to ascertain how new viruses can enter the varroa-bee ecosystem. The fact that it is open to new viruses is troubling, particularly given the recent arrival of varroa in Australia, which is home to a diverse picornavirus community (43) that could be the source of future varroa-transmitted diseases.

One limitation of phylogeographic studies is that sampling biases can affect outcomes since reconstructions of ancestral states are sensitive to the number of samples from a particular geographic region (48). To minimise the potential impact of sampling bias, we attempted to get as broad a range of specimens as possible, including those from the present collection and data publicly available at NCBI. In the case of DWV-A this approach yielded a large number of sequences with good geographic coverage. The number of DWV-B sequences was considerably lower, making it impossible to infer its geographic origins with certainty. However, the reconstructed location of the most ancestral lineages in our data set appears in the Middle East, and an Asian origin is unlikely. Additional sampling of DWV-B, possibly from publicly available data could resolve this uncertainty.

Our study illustrates the dynamic interaction between virus, vector and host. Although the original host switch of *V. destrucor* took place in the mid-20th century, (co-)evolutionary dynamics continue in this system with the acquisition of novel viruses and novel vectors (14, 15). Our findings raise interesting questions about how viruses interact with both varroa and host gene expression, and how viruses evolve to increase their fitness on novel hosts (see for example (21)). The dynamism of the bee-varroa-virus system can make it an excellent model for viral emergence and evolution, as well as broadly applicable strategies for viral control.

## Data Accessibility

Raw sequence data are available under DDBJ/ NCBI BioProject PRJDB14940.

## Acknowledgements

We are grateful to all our generous material providers who supported this work by collecting fresh mites in their apiary or digging in their freezers: Lilia I. De Guzman and Mandy Frake (Brazil and Louisiana, U.S.A); Alison McAfee, Leonard Foster, Heather Higo (British Columbia, Canada); Alina Varaldi and the ROMAPIS beekeepers (Romania); Jevrosima Stevanovic (Serbia); Vivian Wepfer (Switzerland); Peter Rosenkranz (Germany); USDA APHIS (U.S.A); Henriette Rasolofoarivao (Madagascar); Tjeerd Blacquiere (Netherlands); Junichi Takahashi (Japan and New Zealand); and finally the Onna Village Office with the Honey Coral Project (Okinawa). We warmly thank Miyuki Suenaga for technical help with the RNA extraction and library preparation. Library sequencing was performed with the support of the OIST sequencing center. We thank the OIST Scientific Computing and Data Analysis Section for providing HPC to carry data processing. A.S.M. was supported by a Future Fellowship from the Australian Research Council (FT160100178) and a Kakenhi Grant-in-Aid for Scientific Research from the JSPS (18H02216). M.A.T.’s research was supported by a postdoctoral fellowship from the Japan Society for Promotion of Science (JSPS) (P19723) and Kakenhi Grant-in-Aid (19F19723).

